# The neural computations for stimulus presence and modal identity diverge along a shared circuit

**DOI:** 10.1101/2020.01.09.900563

**Authors:** David A. Tovar, Jean-Paul Noel, Yumiko Ishizawa, Shaun R. Patel, Emad N. Eskandar, Mark T. Wallace

**Affiliations:** Vanderbilt Brain Institute, Vanderbilt University, Nashville, USA; Department of Anesthesia, Critical Care & Pain Medicine, Massachusetts General Hospital, Harvard Medical School, Boston, USA; Department of Neurology, Massachusetts General Hospital, Harvard Medical School, Boston, USA; Leo M. Davidoff Department of Neurological Surgery, Albert Einstein College of Medicine, New York, USA; Department of Hearing and Speech, Vanderbilt University Medical School, Nashville, USA; Department of Psychology, Vanderbilt University, Nashville, USA; Department of Psychiatry and Behavioral Sciences, Vanderbilt Medical School, Nashville, USA

**Keywords:** Cortical Circuits, Multivariate Pattern Analysis, Machine Learning, Multisensory, Audition, Touch

## Abstract

The brain is comprised of neural circuits that are able to flexibly represent the complexity of the external world. In accomplishing this feat, one of the first attributes the brain must code for is whether a stimulus is present and subsequently what sensory information that stimulus contains. One of the core characteristics of that information is which sensory modality(ies) are being represented. How information regarding both the presence and modal identity of a given stimulus is represented and transformed within the brain remains poorly understood. In this study, we investigated how the brain represents the presence and modal identity of a given stimulus while tactile, audio, and audio-tactile stimuli were passively presented to non-human primates. We recorded spiking activity from primary somatosensory (S1) and ventral pre-motor (PMv) cortices, two areas known to be instrumental in transforming sensory information into motor commands for action. Using multivariate analyses to decode stimulus presence and identity, we found that information regarding stimulus presence and modal identity were found in both S1 and PMv and extended beyond the duration of significant evoked spiking activity, and that this information followed different time-courses in these two areas. Further, we combined time-generalization decoding with cross-area decoding to demonstrate that while signaling the presence of a stimulus involves a feedforward-feedback coupling between S1-PMv, the processing of modal identity is largely restricted to S1. Together, these results highlight the differing spatiotemporal dynamics of information flow regarding stimulus presence and modal identity in two nodes of an important cortical sensorimotor circuit.

**Significance Statement:** It is unclear how the structure and function of the brain support differing sensory functions, such as detecting the presence of a stimulus in the environment vs. identifying it. Here, we used multivariate decoding methods on monkey neuronal data to track how information regarding stimulus presence and modal identity flow within a sensorimotor circuit. Results demonstrate that while neural patterns in both primary somatosensory (S1) and ventral pre-motor (PMv) cortices can be used to detect and discriminate between stimuli, they follow different time-courses. Importantly, findings suggest that while information regarding the presence of a stimulus flows reciprocally between S1 and PMv, information regarding stimulus identity is largely contained in S1.

## Introduction

Single-unit neurophysiological recordings demonstrate that neural activity within the primary somatosensory area (S1) is monotonically related to stimulus amplitude (Mountcastle et al., 1969). This suggests that a rate code is used to signal the probability of a somatosensory stimulus being present in the environment (Ahissar et al., 2000). Beyond this first cortical area, however, neurons show a variety of response patterns to different stimulus features. For example, some neurons show increasing spiking activity with increasing stimulus frequency, whereas others show the opposite relationship (Salinas et al., 2000). Furthermore, non-linear computations may effectively help filter which information is propagated forward in the cortical hierarchy to solve discrimination problems (Romo & de Lafuente, 2012). Thus, the computational principles that appear best suited for stimulus detection are unlikely to be those best suited for stimulus discrimination. It is currently unclear how brain circuits support these various aspects of processing a sensory stimulus, and how the same brain regions differ in this regard.

Arguably, understanding the mechanistic bases of how the brain signals the presence and the identity of a stimulus has been challenging partly due to the widespread use of univariate techniques and the heavy focus on characterizing the responses of single neurons. However, it is increasingly common to record multiple neurons concurrently across areas, and using multivariate frameworks, uncover neural codes (i.e., response patterns) that are present at the population level (Jonas & Kording, 2017). In addition to understanding the basic characteristics of neural activity of specific neurons and within specific areas, multivariate analyses are able to further probe the manner by which distinct modules communicate with one another, and thus how information is propagated and transformed within the brain (Kumar et al., 2010; Stringer et al., 2019). With large-scale simultaneous multi-area recordings becoming commonplace (Jun et al., 2017; Steinmetz et al., 2018), these analyses are becoming increasingly important tools (Buzsaki, 2004; Stevenson & Kording, 2011).

In the current study, we sought to track information flow relating to the presence and modal identity of a stimulus by examining global neural patterns using multivariate pattern analysis. We simultaneously recorded neuronal activity from two intermediate stages along the hierarchy from sensory input to motor output – primary somatosensory (S1) and ventral pre-motor (PMv) cortex. These areas are two key nodes in a well-established circuit for tactile detection and discrimination (Romo et al., 2004; de Lafuente & Romo, 2005, 2006). In addition to its role in somatosensory function, the PMv cortex is known to be important in auditory discrimination (Lemus et al., 2009) and also possesses multisensory audio-tactile neurons (Graziano et al., 1997). Hence, recording simultaneously from these two areas provides the opportunity to not only examine how information flows between S1 and PMv to support tactile stimulus detection, but also to examine information encoding and flow in the context of determining stimulus modal identity (i.e., auditory, tactile, audio-tactile).

To address this question, tactile, auditory, and audio-tactile stimulation was passively delivered to rhesus monkeys, and neural signals related to the presence and/or modal identity of the stimulus were decoded using multivariate methods (Edelman et al., 1998; Haxby et al., 2001; Kriegeskorte & Kievit, 2013; Goddard et al., 2017). In addition to training and testing within neural areas and at similar time-points, we dissociate these time-periods (time-generalization technique; King & Dehaene, 2014), as well as train and test neural decoders across brain regions. The novel joint application of the time-generalization technique and cross-area decoding allows the tracking of information transfer between S1 and PMv. This combination highlights strikingly different spatiotemporal dynamics in the transfer of information related to the presence vs. modal identity of the stimulus.

## Methods

### Animal Model

Two adult male monkeys (*Macaca mulatta*, 10 –12 kg; Monkey E and Monkey H) were used. Animals were handled according to the institutional standards of the National Institutes of Health (NIH) and protocols were approved by the institutional animal care and use committee at Massachusetts General Hospital.

### Surgical Procedures

A titanium head post and a vascular access port in the internal jugular vein (Model CP6; Access Technologies) were surgically implanted on each of the two animals. Once the animals learned the behavioral task (see below), a craniotomy was performed and extracellular microelectrode arrays (Floating Microelectrode Arrays; MicroProbes) were implanted into S1 and PMv by following landmarks on the cortical surface and stereotaxic coordinates (Fig 1A). Each array (1.95×2.50 mm) contained 16 platinum–iridium recording microelectrodes (0.5 MΩ, 1.5– 4.5 mm staggered length) separated by 400 µm. Monkey E had two arrays in S1 and another two in PMv (total of 32 electrodes in each area, all in the left hemisphere). Implantation for Monkey H was identical to that of Monkey E, with the exception that all electrodes were implanted in the right hemisphere. The recording experiments were performed after 2 weeks of recovery following the array surgery. All experiments were conducted in a radio frequency-shielded recording enclosure.

**Figure 1.**
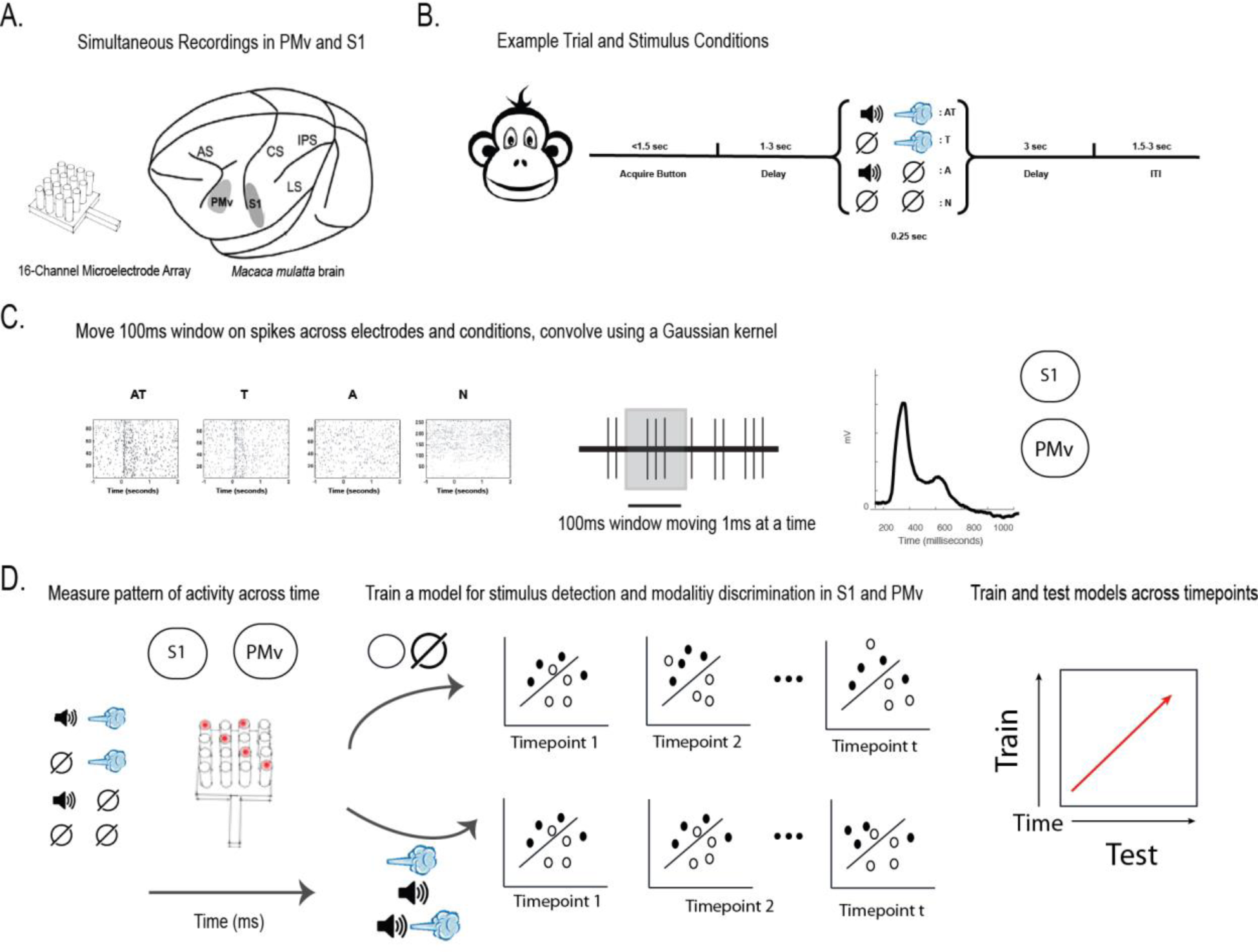
Experiment Schematic. **(A)** Neural recordings were effectuated via 16-electrodes platinum–iridium arrays implanted in S1 and PMv. **(B)** Animals were trained to initiate trials via button press, which following a delay would evoke one of three sensory stimulus sets (audio, tactile, or audio-tactile), or a catch trial with no sensory stimulation delivered. **(C)** Raster plots of an example session in S1 and the average S1 response to tactile stimulation after convolving spike trains with a box-car 100ms in length and moving in 1ms steps. **(D)** Multivariate classifiers (Linear Discriminant Analysis, LDA) were trained on each time-point to differentiate either between the absence and presence of sensory stimuli (regardless of the nature of the stimuli; detection), or to discriminate between sensory modalities (audio, tactile, or audio-tactile; discrimination).

### Materials and Apparatus

Three different types of sensory stimulation were given: audio-alone, tactile-alone, and a combined audio-tactile multisensory conditions. The tactile stimuli were air puffs of 250ms duration delivered at 12 psi to the lower part of the face contralateral to the recording hemisphere. This tactile stimulus was delivered via a computer-controlled regulator with a solenoid valve (AirStim; San Diego Instruments). The eye area was avoided from the puff stimulation. Auditory stimuli were pure tones (4000 Hz at 80 dB SPL) lasting 250ms. These tones were generated by a computer and delivered using two speakers 40 cm from the animal. White noise (50 dB SPL) was applied throughout the trial to mask the air puff and mechanical noises. Audio-tactile stimulation was the synchronous administration of the auditory and tactile stimuli described above. All of the stimulus sets were presented randomly to the animal throughout the recording session.

### Experimental Procedure

After a start tone (1000 Hz, 100 ms), the animals were required to initiate each trial by holding the button located in front of the primate chair using the hand ipsilateral to the recording hemisphere. Animals were required to hold the button until the end of a trial, which was indicated by a liquid reward 3 seconds after stimuli onset (Fig 1B). The monkeys were trained to perform a correct response in >90% of the trials consistently for longer than ∼1.5 h. One of the three sensory stimulus sets (audio, tactile, or audio-tactile), or a catch trial with no sensory stimulation, was delivered to the animal during the trial at a random delay. Each condition was equally likely to be presented.

### Single-Unit Activity, Recording and Preprocessing

Neural activity was recorded continuously and simultaneously from S1 and PMv. Analog data was amplified, band-pass filtered between 0.5 and 8 kHz, and sampled at 40 kHz (OmniPlex; Plexon). Spiking activity was obtained by high-pass filtering at 300kHz and applying a minimum threshold of 3 standard deviations in order to exclude background noise from the raw voltage traces on each channel. Subsequently, action potentials were sorted using waveform principal component analysis (Offline Sorter; Plexon) and binned into 1 ms bins, effectively rendering the sampling rate of 1 kHz. Spike time-stamps were convolved with a 100ms long box-car window and moved in 1 ms steps (Fig 1C). Time-courses were then baseline-corrected by subtracting their pre-stimulus activity (−200 ms to 0 ms post-stimulus onset). This dataset has been previously reported in Ishizawa et al., 2016, and Noel et al., 2019.

### Multivariate Pattern Analysis

Our aim here was to track message passing and information transformation within the cortex and hence focus on multivariate decoding techniques. Following data preprocessing, we used CoSMoMVPA (Oosterhof, et al., 2016) to decode stimulus presence vs. absence, as well as the sensory modality of the stimuli presented. Linear discriminant analysis (LDA; Duda et al., 2001) classifiers were trained and tested in 1ms increments using 4-fold cross validation. In this procedure, trials are randomly assigned to one of four subsets. Three of the four subsets (75% of the data) are pooled together to train the classifier and then decoding accuracy is tested on the remaining subset (25% of the data). This procedure is repeated a total of four times, such that each of the subsets is tested once. Decoding results are reported in percent correct of classifications at each time point in the time series ranging from −100ms to 1000ms relative to stimulus onset. This analysis was conducted independently for each recording session (n=18), distinction of interest (stimulus presence and stimulus modality), as well as within and across brain areas (S1 and PMv). Mean and standard error were then calculated across recording sessions at each time point (Fig 1D).

Regarding statistical analyses, each time point was tested for the null and alternative hypotheses using Bayes’ factors. The null hypothesis indicates that there is no information regarding the presence or absence of stimuli for stimulus detection and no information regarding the type of modality for modality discrimination. Thus, the null hypothesis would be the decoder guessing at chance, which would be 50.0% decoding accuracy for stimulus presence, and 33.3% decoding accuracy for modality discrimination. We then calculated the probability of the alternative hypothesis in relation to the null hypothesis. A Bayes’ factor greater than 3 indicates substantial evidence for the alternative hypothesis, anything between 3 and 1/3 indicates insufficient evidence, and values less than 1/3 indicate evidence for the null hypothesis (Jeffreys, 1961; Wetzels et al., 2010). Substantial evidence for the alternative hypothesis indicates that the brain state contains meaningful information that the classifier can utilize to identify the correct trial condition for stimulus presence (stimulus present or absent) or stimulus identity (audio, tactile, or audio-tactile). Furthermore, Bayes’ factors provide an added advantage over Frequentist inference: in addition to rejecting the null hypothesis, this framework can also provide support for either the null hypothesis or to determine that the data is insensitive. For both stimulus presence and modality discrimination, trials were balanced across conditions, as imbalance among class types can have the unwanted effect of biasing the classifier toward the class with more trials (Grootswagers et al, 2017).

### Time Generalization Within Areas and Across Areas

To probe the dynamics of the available information used by the classifier to decode presence or absence of stimuli regardless of modality, as well as the modality of the stimuli presented, we used a time generalization decoding technique (Carlson et al., 2011; King & Dehaene, 2014; Fig 1D). In this analysis, the classifier was trained on the same decoding distinctions as before (presence and identity of sensory stimuli). However, to investigate how well neural data from one timepoint generalizes to all others, the classifier is trained on a particular timepoint within the time series (i.e., −100 to 1000 ms post-stimuli onset) and then tested with data from every timepoint in the time series. This procedure was repeated for every timepoint and concatenated to create 1100 x 1100 matrix containing every possible combination of training and testing timepoints. The diagonal along the matrix represents times in which training and testing were performed within the same timepoint. Lastly, we performed a similar time generalization analysis across areas in order to investigate how well different timepoints in one area can decode information at different timepoints in the other area – putatively indicating the flow of information from one area at one time-point, to another area at a different time-point. We trained across all timepoints in S1 and tested on PMv and then performed training on PMv and tested on S1. Since PMv and S1 had an unequal number of single units captured (S1 = 9.6 +/− 4.5, PMv = 5.7 +/− 2.3) we randomly subsampled from the area with more single units isolated. To eliminate potential sampling bias, we performed the cross-area time generalization analyses ten times, with different randomly subsample single units. The mean decoding results across the ten iterations was then computed for each recording session.

## Results

### Multivariate Decoding Allows Tracking Stimuli Presence and Identity Over Long Periods

Both S1 and PMv showed evoked responses during the presentation of sensory stimuli. We used Bayes factors at each timepoint to assess whether the evoked responses diverged significantly from baseline activity. For the univariate analysis, when averaging responses over modalities, S1 showed a strong evoked response, showing substantial evidence for the alternative hypothesis (defined as Bayes factor [BF] >3) at two-time periods, from 19-184 ms and from 309-388 ms post-stimulus onset. PMv showed a later response, from 141-422 ms post-stimulus onset. When looking at evoked responses to specific modalities, S1 responds to tactile stimulation for the period between 36-184 ms post stimulus onset, to auditory stimuli from 21-102 ms post-stimulus onset, and to audiotactile stimulation from 19-197 ms and 328-414 ms post-stimulus onset. Responses of PMv to sensory stimuli are not as robust, but there is clear evidence for evoked responses to tactile stimuli from 131-157 ms and from 212m-448 ms post-stimulus onset, to auditory stimuli from 182-405 ms post-stimulus onset, and to combined audiotactile stimuli from 166-381 ms post-stimulus onset.

Using time-resolved LDA, we were able to decode the presence (vs. absence) of stimuli in S1 and PMv (Fig 2C). Onset decoding latencies, defined as the first timepoint of at least 20 ms of sustained significant decoding above chance (see Carlson et al., 2013) were found for S1 beginning 36 ms post-stimulus onset (Fig 2C, purple) and for PMv beginning at 58ms post-stimulus onset (Fig 2C, green). Maximum decoding performance was reached at 183 ms post-stimulus onset for S1 and at 222 ms post-stimulus onset for PMv. For both S1 and PMv, decoding remains significantly above chance for periods extending beyond 1000 ms post stimulus onset. This observation highlights the utility of indexing not only the activity of single neurons via traditional univariate approaches, but also in examining the responses of neuronal populations via multivariate decoding. For example, the average firing rate produces a strong transient response to tactile stimulation followed by a sustained response in S1, which return to baseline within approximately 500ms. In contrast, it was possible to decode the presence of a stimulus in S1 for a period at least twice as long (~ 1000ms) using multivariate approaches.

**Figure 2.**
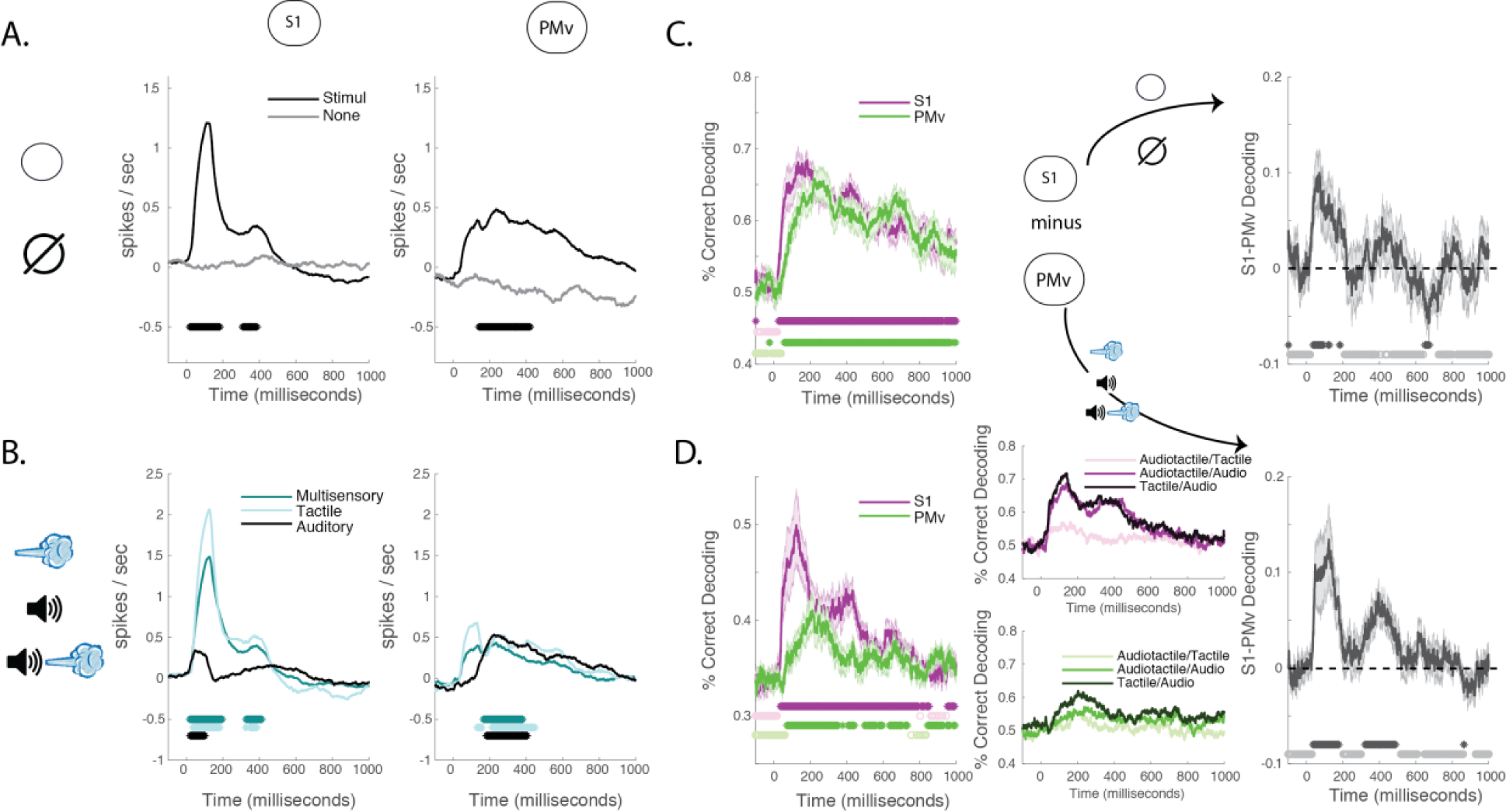
Univariate and Multivariate Responses to Sensory Stimulation. **(A)** Both S1 and PMv show evoked responses during the presentation of sensory stimuli. **(B)** S1 responds to tactile stimulation, while also responding to audio-tactile stimulation, but less to auditory stimuli alone. PMv does not show as clear evoked responses to sensory stimuli as primary somatosensory area does but shows less disparity in evoked responses across stimuli types. **(C)** LDA classified above chance either the presence or absence of sensory stimulation starting 36ms and 58ms for S1 and PMv respectively post-stimuli onset, and lasting 1s, well beyond the time-period where univariate responses are apparent. As illustrated by the difference in correct decoding between S1 and PMv, information regarding stimulus detection was present first in S1, then was present in both S1 and PMv, and finally was stronger in PMv than S1. **(D)** Discrimination of sensory modalities was also correctly decoded by LDA, with modal identity being clearer in S1 than PMv, particularly at early latency post-stimulus onset, and between approximately 200 and 400ms post-stimulus onset. Asterisks indicate significant decoding above chance, using Bayes’ factors (Bayes’ Threshold >3).

We next used Bayes’ factors to look at the time-resolved differences in the decoding of stimulus presence between S1 and PMv. Results demonstrate significant evidence supporting the alternative hypothesis (BF>3), suggesting a differential time-course during which stimulus presence information is available in S1 and PMv (Fig 2C, black curve). Beginning at 40 ms and extending up until 186 ms, decoding was better in S1 than PMv, consistent with the earlier decoding onset found in the primary sensory area. Following 186 ms, evidence is stronger for the null hypothesis (BF<1/3) up until 651 ms post-stimulus onset. Following 651 ms, evidence for the alternative hypothesis is once again supported, but this time in PMv. These findings suggest that information regarding stimulus presence may be transferred between S1 (first) and PMv (later).

We then applied the same approach to determining when the modality (i.e., A, T, AT) of the stimulus could be decoded from the neural signals in S1 and PMv. Results suggested above chance decoding (i.e., >33.3%) starting 37 ms post-stimulus onset for S1 (Fig 2D, purple), and starting 70 ms post-stimulus onset for PMv (Fig. 2D, green). A maximum modality decoding performance of 50.0% was reached at 125 ms post-stimulus onset for S1 and a maximum modality decoding performance of 41.0% was reached at 213 ms post-stimulus onset for PMv. As shown by the difference in decoding performance within S1 and PMv (Fig 2D, black curve), decoding accuracy was significantly higher for S1 relative to PMv for two sustained periods - between 40-74ms post-stimulus onset, as well as between 348-470ms post-stimulus onset. Collectively, these results suggest that the computations underlying the detection of a stimulus and the identification of stimulus modality evolve over differing temporal epochs in S1 and PMv. More specifically, while information regarding detection appears later in PMv as compared to S1, and thus leading to a single time-period where stimulus presence is more readily decoded in S1 than PMv, information regarding stimulus modality is more readily decoded in S1 over PMv over both an early and late temporal epoch.

### Information Regarding the Presence and Modal identity of Stimuli Follow Different Dynamics in S1 and PMv

To further explore how information dynamics regarding the encoding of the presence or modal identity of a stimulus varies across S1 and PMv, we used a time generalization approach (Carlson et al., 2011; King & Dehaene, 2014) where a classifier is trained at one timepoint and then tested across the remaining timepoints. Specifically, it probes how information at a given timepoint generalizes to information throughout the time series to understand whether the information is increasing, decreasing, or re-emerging at later times. In the present study we leveraged the fact that decoding performance is better when training on a low signal-to-noise ratio (SNR) and testing on a high SNR (Fig 3A, van den Hurk & Op de Beeck, 2019) to quantify the degree to which information at a particular time is changing (Fig 3B). Given that training timepoints are plotted along the y-dimension and testing times are plotted along the x-dimension in a time generalization matrix, if information at a particular timepoint increases during the time-course, this will appear as an off-diagonal shift in the horizontal (rightward) direction. Conversely if information at a timepoint decreases during the time course, this will appear as an off-diagonal shift in the vertical (upward) direction. To calculate the overall direction of information change (horizontal or vertical off-diagonal) we subtracted the vertical off-diagonal from the horizontal off-diagonal.

**Figure 3.**
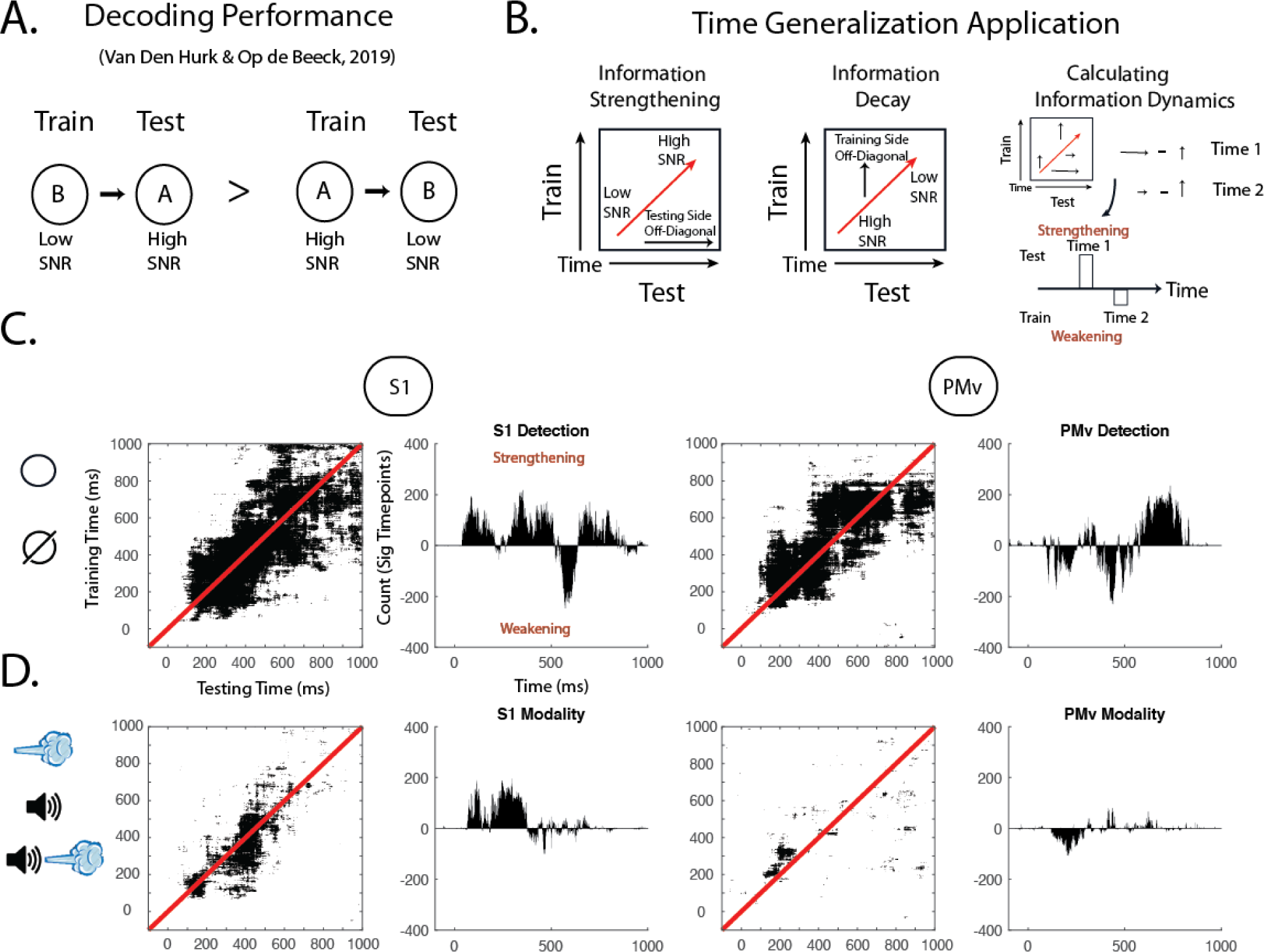
Time generalization results. **(A)** Decoding accuracy is better when training on low signal-to-noise (SNR) and testing on higher SNR, than when training on high SNR and testing on low SNR. **(B)** In conjunction with the time-generalization technique, this observation can be leveraged to estimate whether a particular information state is strengthening (i.e., becoming more discriminant with time) or weakening within a cortical area. **(C)** Time generalization plots for decoding stimulus presence in S1 (left) and PMv (right). Second and fourth columns show the difference in significant time-points over which decoding generalizes along the training and testing axis – positive counts indicate information strengthening, while negative counts indicate information weakening. **(D)** Follows the same convention as **(C)**, for modal identity as opposed to stimulus presence.

Regarding the decoding of stimulus presence, we calculated all of the times where there was very strong evidence (BF>30) for the alternative hypothesis to capture the most prominent information states (Fig 3C). The plots showed that decoding onset and off-diagonal spread across training-testing time periods varied between S1 and PMv for information regarding stimulus presence (see Fig 3C, first and third columns). This difference becomes apparent in the horizontal-vertical off diagonal difference histograms (see Fig 3C, second and fourth columns). In S1, the information states strengthen from 39-207ms and then again from 239-555 ms, after which information weakens until 638 ms and then ends with a final wave of strengthening. On the other hand, PMv shows the opposite pattern, with information initially weakening beginning at 93 ms, then briefly strengthening at 261 ms, weakening again at 351 ms, and showing a final wave of strengthening beginning at 497 ms. Overall, these results highlight a general pattern in which these are opposing temporal dynamics to information strengthening and weakening for S1 and PMv (see Supplemental Fig 1), potentially implying that information is flowing back and forth between these two areas. Furthermore, whereas information regarding stimulus presence demonstrated a greater strengthening pattern in the initial response epoch for S1, PMv demonstrated a greater strengthening pattern in the late time period (after 500ms).

In contrast to the patterns seen for information regarding stimulus presence, the dynamics for information about modal identity in S1 and PMv show consistent strengthening and weakening, respectively. In S1, information regarding modal identity begins to increase at 65 ms and continues to increases until 387 ms. In contrast, in PMv information regarding modal identity shows a weakening pattern beginning at 90 ms and continues to weaken until 299 ms. In sum, the difference in dynamics regarding information pertaining to stimulus presence and modal identity strongly suggest differences in how this information is processed and shared between S1 and PMv. An additional finding that is illustrated by the on-diagonal analyses is that overall information regarding modal identity is short-lived as compared to information regarding stimulus presence.

### Cross-Region Decoding Reveals Feedforward Presence Information and Feedback Identity Information

To more directly track shared information between S1 and PMv, we trained classifiers on neural data collected from one region and tested on another, while also performing time-generalization (Carlson et al., 2011; King & Dehaene, 2014). For example, and as illustrated in Fig 4A, we can use this analysis to train on S1 and test on PMv to examine for potential significant horizontal off-diagonals (i.e., in the future along the testing dimension). Such a result would suggest that S1 shares common information that is present at a later time in PMv (see Fig 4A for other examples).

**Figure 4.**
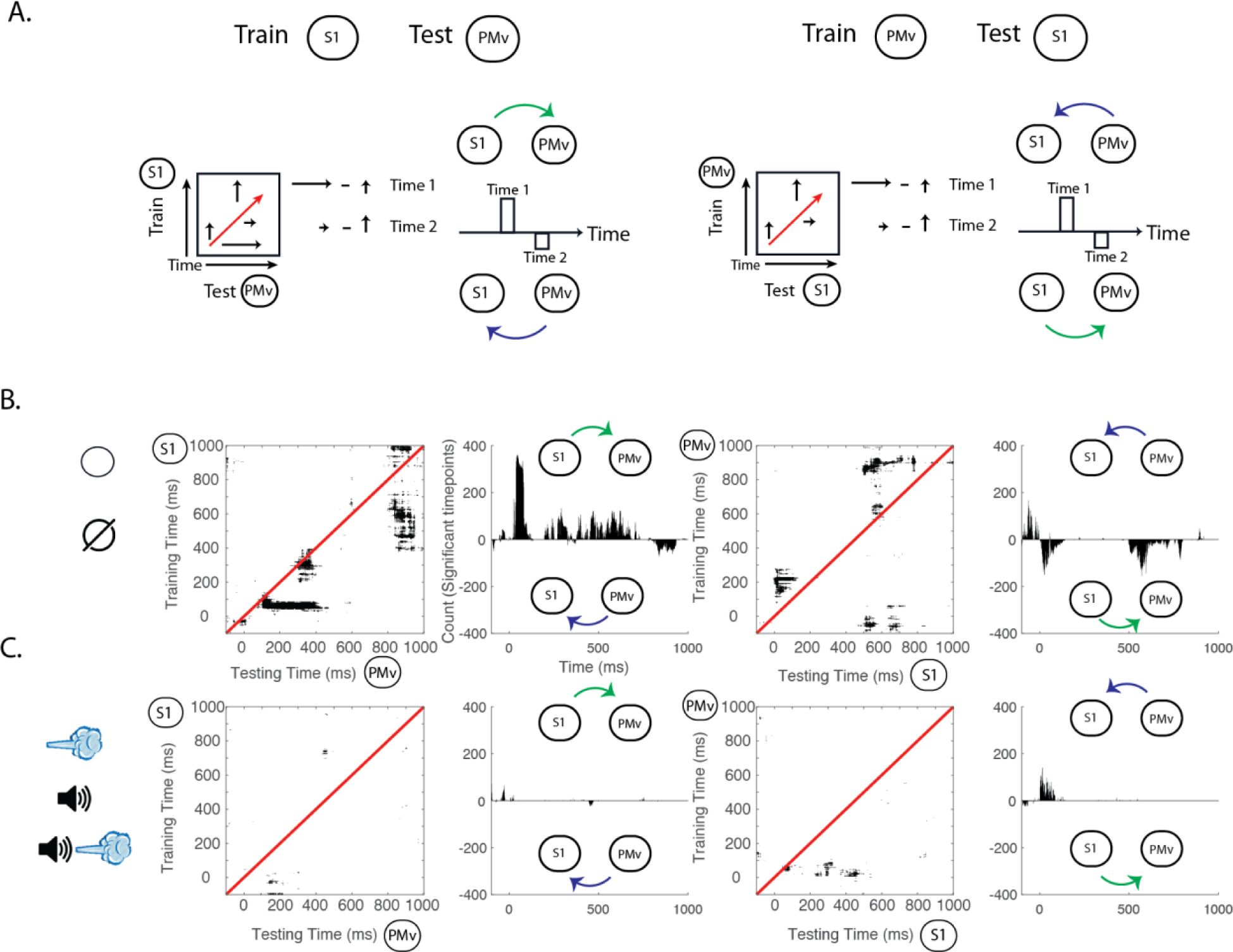
Cross-area time series decoding and time generalization. **(A)** Schematic showing two possible direction in which information can generalize. For training on S1 and testing on PMv. If information generalizes from S1 to PMv, a horizontal off-diagonal will be seen when training. On the other hand, an off-diagonal is vertical (i.e., later training periods can decode earlier ones), information is generalized in the direction of PMv to S1. **(B)** Cross-area off-diagonal examination for stimuli presence decoding in S1 and PMv. Results show a clear horizontal off diagonal when training in S1 and decoding in PMv. Training on PMv and decoding in S1 demonstrates a vertical off-diagonal **(C)** Cross area decoding of the identity of sensory stimuli. Training LDA in S1 does not afford the possibility of decoding sensory modality in PMv. Contrarily, training on PMv shows off-diagonal decoding along the testing dimension, suggesting information generalizes from PMv to S1.

In decoding stimulus presence we found that training and testing across S1 and PMv along the same time points did not yield any periods of time with sustained significant decoding accuracy using a criterion of substantial evidence (BF>3) (Fig 4B **on-diagonal**). Such a result suggests that S1 and PMv do not contain common information regarding the presence of a stimulus during the same time period, although both do contain information regarding stimulus presence (Fig 2C). The fact that the within-area decoding is successful, but across area decoding is not, suggests that the codes for stimulus presence within S1 and PMv are likely of different format.

To better explore whether the lack of simultaneous shared information was due to a transformation of information from one area to another, we inspected the off-diagonals in the time generalization matrices. As shown in Fig 4B **off-diagonal**, results indicate that beginning at approximately 38ms and extending to 100ms post-stimulus onset, information regarding stimulus presence in S1 significantly generalizes to PMv for the time period spanning between ~100-442 ms post-stimulus onset This result shows that information pertaining to stimulus presence in PMv at this later interval is similar to that seen earlier (38ms to 100ms) in S1. Training on PMv and attempting to decode within S1 across different time periods also yielded significant vertical off-diagonals (i.e., along the training dimension), beginning at 15 ms post-stimulus onset and extending forward to 97-287 ms. (Fig 4B). Thus, whether training in S1 or PMv, information regarding stimulus presence generalized in the direction from S1 to PMv. The fact that training in PMv and decoding in S1 yielded a more restricted time-period of generalization than within area decoding in S1 and PMv (Fig 2C) could imply that while information regarding stimulus presence in PMv was initially of the same format as that in S1, it is subsequently transformed in such a way that the new format could not be generalized back to S1.

Regarding the discrimination of modal identity, just as for the decoding of stimulus presence, we found that training and testing along the same time points did not yield any time periods with significant and sustained decoding accuracy (Fig 4C). Extending sensory modality classifiers trained in S1 to PMv along the time generalization matrices did not demonstrate any time periods of successful classification (Fig 4C). On the other hand, when we trained in PMv and tested on S1, at 10 ms post-stimulus onset there was a higher than chance decoding accuracy in S1 along an array of time-points in the future. Thus, very early patterns of activity supporting the classification of modal identity in PMv are later found in S1. Thus, unlike stimulus presence, it appears that modal identity information generalizes in the direction of PMv to S1.

## Discussion

Simultaneous recordings of spiking activity across distinct nodes of a canonical sensorimotor circuit allowed us to study how information is shared and transformed between these areas. We recorded from arrays of electrodes placed in S1 and PMv – two areas known to be instrumental in transforming sensory information from different modalities (tactile, as well as auditory) into motor commands for action (Romo et al., 2004; de Lafuente & Romo, 2005, 2006; Graziano et al., 1997; Noel et al., 2019). Specifically, we were interested in how information regarding stimulus presence and modal identity flowed and was altered between S1 and PMv and used different multivariate analyses to examine this question. The principal findings of the study are: 1) for decoding the presence of a stimulus, decoder performance fluctuated in a reciprocal manner between S1 and PMv for the interval up to 1 second after stimulus presentation, while decoding of modal identity was consistently higher in S1 than in PMv, 2) using time generalization, information regarding stimulus presence showed oscillatory strengthening and weakening dynamics in both S1 and PMv, while information regarding modal identity exhibited steady strengthening in S1 and weakening in PMv, 3) using cross-area time generalization, information regarding stimulus presence generalized between S1 and PMv, offset in time in the direction of S1 to PMv, while modal identity information only generalized weakly from PMv to S1. Together, these results highlight the different dynamics for the flow of information regarding stimulus presence and modal identity in two nodes of an important cortical sensorimotor circuit.

The findings fit within a larger and longstanding debate in neuroscience regarding whether sensory modality information is preserved as it ascends the processing hierarchy, or if that information ultimately transitions into an amodal format (Machery, 2016). Evidence for modality-specific information being preserved at high levels of representation comes from mental imagery, priming, and dreaming studies which show recruitment of sensory specific areas in the brain that are similar to their respective perceptual counterparts (Caramazza & Mahon, 2003; Horikawa, Tamaki, Miyawaki, & Kamitani, 2013; Ishai, Haxby, & Ungerleider, 2002). On the other hand, evidence for amodal representations include task-specific recruitment of common brain areas for representations such as magnitude and numerosity regardless of sensory modality (Piazza, Mechelli, Price, & Butterworth, 2006). Additionally, blind patients who hear sounds corresponding to objects viewed by sighted individuals shows similar brain activations (van den Hurk, Van Baelen, & Op de Beeck, 2017). Our cross-area time generalization results provide evidence, that at least in the context of the passive delivery of stimuli studied here, as this perceptual information is hierarchically processed in the brain and transferred from sensory regions (S1) to regions closer to the motor circuitry (PMv), the representations become more amodal. Specifically, we found that information regarding stimulus presence in S1 generalized to PMv, but that information regarding modal identity only weakly generalized in the opposite direction from PMv to S1. However, it is important to note that our recordings were limited to S1 and PMv, and thus we cannot claim that modal identity is not preserved in other parts of the sensorimotor (or beyond sensorimotor) hierarchy. Ostensibly, the modal identity information transfer from PMv to S1 may represent the contribution of other nodes to modal identity that PMv is propagating backwards to S1.

In addition to what sensory information is transferred between brain areas, an equally important question is how sensory information is transformed as it ascends the sensory hierarchy. One important manner in which information can be transformed is through recurrent feedback (O’Connell, Dockree, & Kelly, 2012). Notably, visual studies have found feedforward responses predominate during the first 200 ms of visual processing (Dehaene & Changeux, 2011; Thorpe, Fize, & Marlot, 1996) and thereafter recurrent process derived from temporal, parietal, and prefrontal cortices shape and ultimately transform the nature of these visual signals (Gold & Shadlen, 2007; Gwilliams & King, 2019; Kar, Kubilius, Schmidt, Issa, & DiCarlo, 2019; Lamme & Roelfsema, 2000). In our study, we found evidence of a recurrent process for encoding information regarding the presence of a stimulus in S1 and PMv. Notably, when we compared results from the univariate and multivariate analyses, we found the decoding results to reveal that information pertaining to the presence of a stimulus was sustained for up to 1000 ms in both S1 and PMv, well beyond what averaged univariate responses revealed. This difference potentially reflects a change in information format from a standard rate code visible to univariate analyses to a code more reliant on sparse spatio-temporal patterns across the population that is only revealed through the application of multivariate methods. Further, information regarding stimulus presence was found to oscillate between strengthening and weakening in S1 and PMv up until 500 ms, after which it shows a steady strengthening (Fig. 3C). This oscillation coincides with an initial information transfer in the first 500 ms between S1 and PMv noted in the cross-area time-generalization results (Fig 4B). Thus, our results suggest that initial information regarding stimulus presence for auditory, tactile, and audiotactile stimuli is first transferred between S1 and PMv and finally transformed for subsequent decision making.

One caveat of the current results lies in the passive nature of the task, in which the monkey was not required to detect or discriminate between stimuli, but rather was only required to acquire the button after a start tone and release once it received a reward (thus maintaining vigilance). In many respects, this makes the results both surprising and compelling, in that there was no behavioral need to make use of the presented sensory information. Prior work in S1 has shown that responses to passive stimuli are depressed and have much more variability than when the animal is participating in an active process (Crochet & Petersen, 2006; Schroeder, Wilson, Radman, Scharfman, & Lakatos, 2010), and work in the visual system has shown that active tasks have longer sustained decoding than passive tasks when viewing identical stimuli (Ritchie, Tovar, & Carlson, 2015). Collectively, this points to the current work representing an important foundation for future studies, as it illustrates the ability of decoding approaches to reveal differences in the dynamics of information flow in a classic sensorimotor cortical circuit – even when the stimuli are not used in the execution of an action. Future work should require animals to detect and/or discriminate between sensory stimuli, in order to examine whether task demands potentially lead to longer periods of information transfer, and if information regarding modal identity is transferred in a discrimination task dependent upon stimulus identity. Moreover, an active task would allow the establishment of direct links between decoding performance and behavior through the use of distance from the decoding boundary in order to predict metrics such as accuracy and reaction time (Carlson, Ritchie, Kriegeskorte, Durvasula, & Ma, 2014; Ritchie & Carlson, 2016; Ritchie et al., 2015).

In conclusion, we have leveraged the ability to generalize neural activity across both space and time using multivariate techniques in order to garner insights into how information flows and is transformed from low level sensory areas to premotor areas in the brain, where it can be utilized for action. Specifically, we are able to provide empirical support that sensory information from S1 to PMv transitions to an amodal representation in the passive task that was employed. Importantly, our work provides a framework for which future work can explore how sensory information is transferred and transformed beyond just the two brain areas explored in this study and will allow examination of how task demands affect information flow.

## ACKNOWLEDGMENTS

D.A.T. is supported by a by NIGMS of the National Institutes of Health (T32GM007347). J.P.N. was supported by fellowship F31MH112336 by NIMH/NIH. We thank Alex Maier and Adriana Schoenhaut for helpful comments on an earlier version of this manuscript.

## Author’s Contributions

D.A.T., J.P.N., and M.T.W. conceptualized the study. M.T.W., and E.E. supervised the study. Y.I. and S.P. collected the data. Y.I., S.P. and J.P.N., preprocessed the data. D.A.T. performed the main data analyses and created all figures. D.A.T and J.P.N. wrote the initial draft of the paper. D.A.T., J.P.N., Y.I., S.P., E.E., and M.T.W. revised the paper and approved the final version of the manuscript.

## Supplemental Figures

**Figure S1.**
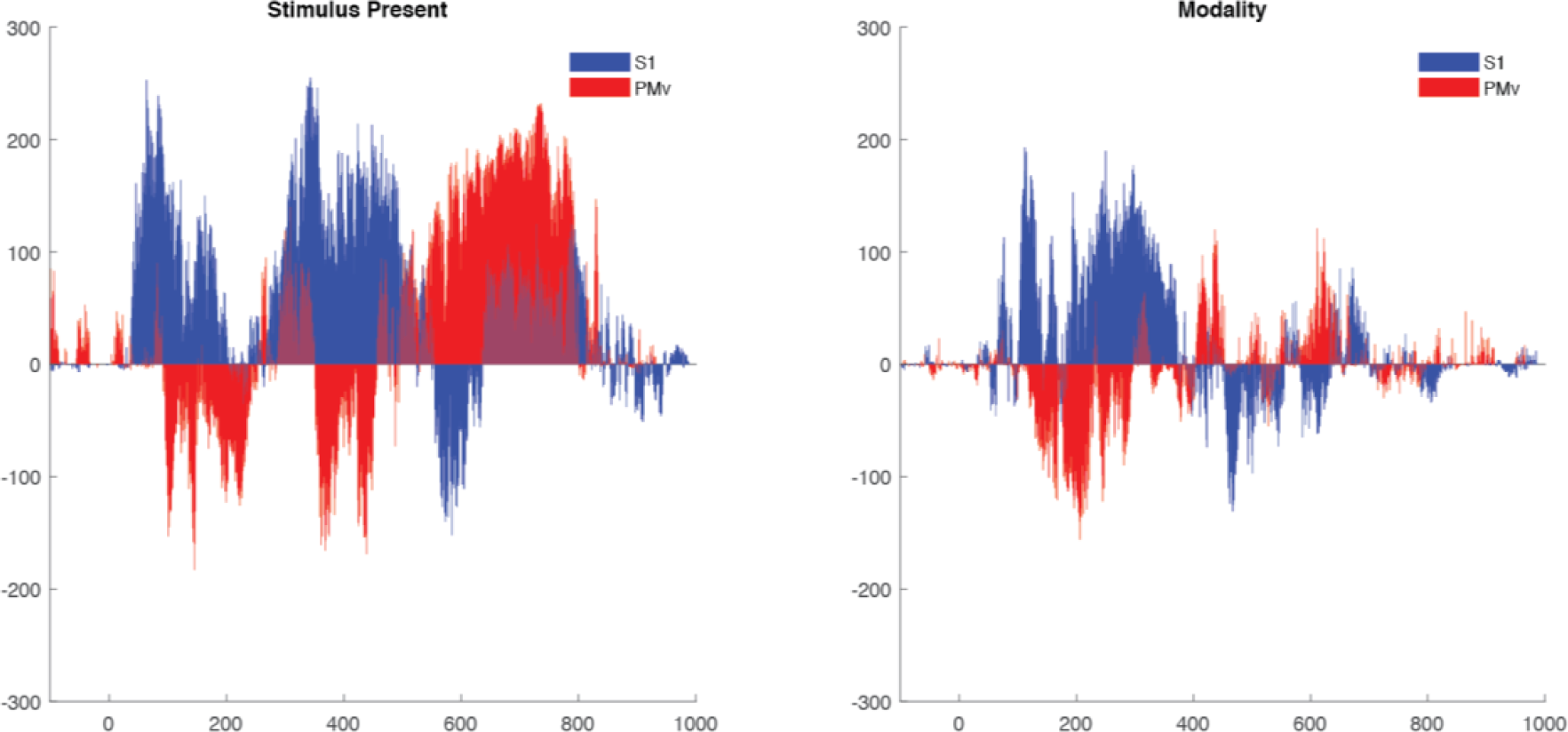
Time Generalization dynamics for stimulus presence and modal identity in S1 and PMv.

**Figure S2.**
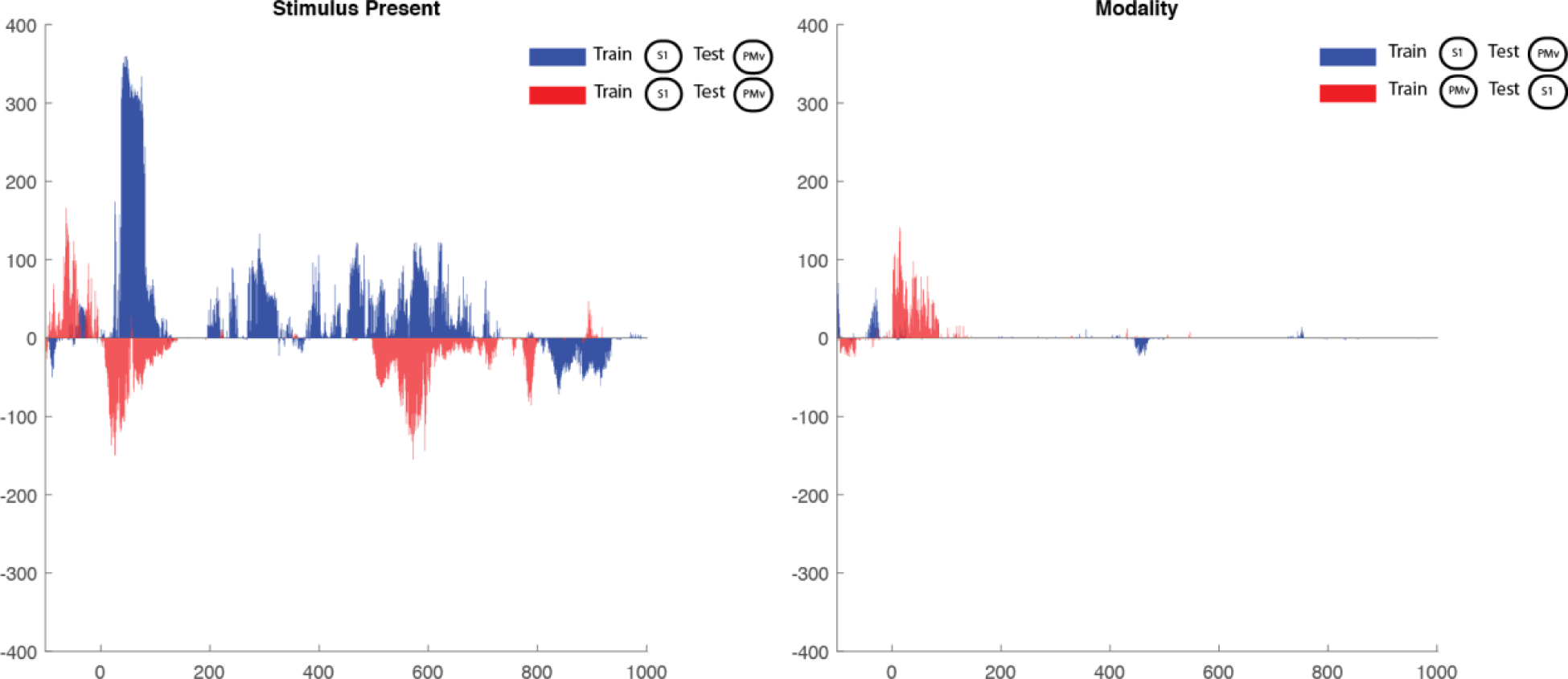
Cross area time generalization dynamics for stimulus presence and modal identity in S1 and PMv, training in S1 and testing in PMv and vice versa.

